# Retrospective varying coefficient association analysis of longitudinal binary traits: application to the identification of genetic loci associated with hypertension

**DOI:** 10.1101/2022.10.31.514543

**Authors:** Gang Xu, Amei Amei, Weimiao Wu, Yunqing Liu, Linchuan Shen, Edwin C. Oh, Zuoheng Wang

## Abstract

Many genetic studies contain rich information on longitudinal phenotypes that require powerful analytical tools for optimal analysis. Genetic analysis of longitudinal data that incorporates temporal variation is important for understanding the genetic architecture and biological variation of complex diseases. Most of the existing methods assume that the contribution of genetic variants is constant over time and fail to capture the dynamic pattern of disease progression. However, the relative influence of genetic variants on complex traits fluctuates over time. In this study, we propose a retrospective varying coefficient mixed model association test, RVMMAT, to detect time-varying genetic effect on longitudinal binary traits. We model dynamic genetic effect using smoothing splines, estimate model parameters by maximizing a double penalized quasi-likelihood function, design a joint test using a Cauchy combination method, and evaluate statistical significance via a retrospective approach to achieve robustness to model misspecification. Through simulations, we illustrated that the retrospective varying-coefficient test was robust to model misspecification under different ascertainment schemes and gained power over the association methods assuming constant genetic effect. We applied RVMMAT to a genome-wide association analysis of longitudinal measure of hypertension in the Multi-Ethnic Study of Atherosclerosis. Pathway analysis identified two important pathways related to G-protein signaling and DNA damage. Our results demonstrated that RVMMAT could detect biologically relevant loci and pathways in a genome scan and provided insight into the genetic architecture of hypertension.

## 1 INTRODUCTION

Genome-wide association studies (GWAS) have successfully identified thousands of susceptible loci underlying human diseases and complex traits. Many epidemiological studies, such as Framingham Heart Study (FHS) and Women’s Health Initiative (WHI), have collected and measured health conditions and phenotypic traits on study participants over the years. Such studies provide rich resources for the investigation of genetic architecture and biological variations of complex disorders. In recent years, more and more genetic studies have exploited genetic data from human biobanks and extracted health information from electronic health record (EHR) data to gain better understanding of the genetic mechanism over the course of complex diseases.

Traditional genetic association analyses on single time point measure fail to capture phenotypic variation over time and may lose statistical power to identify disease-related variants. Thus, genetic studies with longitudinal phenotypes require powerful analytical tools for optimal analysis. Statistical methods that account for dependence structure among observations from the same subject have been developed in GWAS to make full use of longitudinal data, such as mixed effects models (Furlotte, Eskin and Eyheramendy, 2012; Sikorska et al., 2013; Wu et al., 2019), generalized estimating equations (GEEs) (Sitlani et al., 2015; Wu et al., 2019), growth mixture models (Das et al., 2011; Londono et al., 2013), and empirical Bayes models (Meirelles et al., 2013). Most of these methods assume that genetic contribution is constant over time. However, disease development and progression is a complicated process that changes over time. Windows of susceptibility and critical periods across the lifespan exist in disease onset and development. Studies have shown that genetic influence on the trait variations fluctuates with the passage of time (Bryois et al., 2017; Chu, Li and Reimherr, 2016; Gong and Zou, 2012; Liu, Li and Wu, 2014; Wang, Li and Huang, 2008). Modeling time-varying genetic effects is essential to identify and validate causal genetic loci that are associated with time-dependent variation of disease progression.

Varying coefficient models are a class of generalized regression models in which the coefficients are allowed to vary smoothly with the value of other variables (Hastie and Tibshirani, 1993). They are semi-parametric models that explore dynamic pattern in the data to improve model fitting (Fan and Zhang, 2008), reduce model bias by specifying the coefficients as smooth nonparametric functions (Lu and Zhang, 2009), and overcome the “curse of dimensionality” in the nonparametric estimation of multiple regression problems (Eubank et al., 2004). There are several approaches to estimate time-varying coefficients in varying coefficient models, including kernel-local polynomial smoothing (Fan and Zhang, 1999; Hoover et al., 1998; Kauermann and Tutz, 1999; Wu, Chiang and Hoover, 1998), polynomial spline (Huang and Shen, 2004; Huang, Wu and Zhou, 2002, 2004), and smoothing spline (Chiang, Rice and Wu, 2001; Hastie and Tibshirani, 1993; Zhang, 2004). Other statistical methods have been developed to model dependency in longitudinal data, such as GEE (Liang and Zeger, 1986) and Gaussian copula (Joe, 2014). Models allowing for time-varying covariate effects include semiparametric regression with GEE (Lin and Carroll, 2000), and time-varying copula models (Kürüm et al., 2018; Kürüm et al., 2016).

Varying coefficient models have been used for two types of applications in longitudinal GWAS. The first application focuses on feature selection for longitudinal outcomes with ultrahigh-dimensional predictors such as single nucleotide polymorphisms (SNPs). As computational burden is a major concern when handling millions of SNPs simultaneously in a model, feature screening becomes an efficient solution to filter out unimportant SNPs. Feature screening in varying coefficient models has been developed based on conditional Pearson correlation (Liu et al., 2014), extended B-splines (Fan, Ma and Dai, 2014), modified weighted least squares estimation (Chu et al., 2016) and functional regression with group penalty (Marchetti-Bowick et al., 2016) to retain important SNPs associated with continuous and binary traits (Chu et al., 2020; Xia, Yang and Li, 2016). The second application focuses on the detection of time-varying effect of quantitative trait nucleotide, such as functional GWAS (Das et al., 2011; Li et al., 2015; Ning et al., 2017). These methods fit the model at each SNP separately and use likelihood ratio tests to determine statistical significance, thus can be computationally intensive to analyze genome-wide SNPs, especially for binary outcomes. For large GWAS, score tests are popular and computational efficient because they only fit the null model once for all SNPs. However, current score test-based methods commonly treat the effects of genetic variants as constant over time and are not able to capture the dynamic contribution to disease progression.

Motivated by a genome-wide association analysis of longitudinal measure of hypertension in the Multi-Ethnic Study of Atherosclerosis (MESA), we developed a retrospective varying coefficient mixed model association test, RVMMAT, to detect time-varying genetic effect on longitudinal binary traits. We model dynamic genetic effect using smoothing splines, estimate model parameters by maximizing a double penalized quasi-likelihood function, design a joint test using a Cauchy combination method, and evaluate significance of the test via a retrospective approach in which genotypes are treated as random conditional on the phenotype and covariates. Retrospective association tests have been shown to be robust to the trait model misspecification and improve statistical power (Hayeck et al., 2015; Jiang, Mbatchou and McPeek, 2015; Wu et al., 2019; Wu and McPeek, 2018). In RVMMAT, flexible assumptions on the effect function increased power to detect genetic variants associated with dynamic traits. The validity of RVMMAT does not depend on the variance estimation in the trait model due to the tuning parameters in penalty terms. For comparison, we also developed VMMAT, a prospective varying coefficient mixed model association test. We conducted simulation studies to evaluate the type I error and power of RVMMAT and VMMAT, and compared them with the existing association methods. The results demonstrated that the retrospective varying-coefficient test had better control of type I error when the trait model was misspecified, and was robust to various ascertainment schemes. Moreover, the retrospective varying-coefficient test was more powerful than the prospective test. We applied RVMMAT and VMMAT to the genome-wide association analysis of longitudinal measure of hypertension in MESA and identified hypertension-related genetic loci and pathways.

## 2 METHODS

Suppose a binary trait is measured over time on a sample of *n* subjects. We have their genome-wide measures of genetic variation and a set of covariates. The covariates are allowed to be static variables such as sex or dynamic variables such as body weight. Let ***X***_*ij*_ and *Y_ij_*, *i* = 1, …, *n*, *j* = 1, …, *m_i_*, denote the *p*-dimensional covariate vector and the binary trait measured on subject *i* at time *t_ij_*. Here, the measurement time and length are allowed to be different for different subjects. We let ***X*** = (***X***_1,1_, _,***X***_1,*m*_1__,…, ***X***_*n*,1_, …,***X**_n,m_n__*) denote the *N* × *p* covariate matrix and ***Y*** = (*Y*_1,1_, …, *Y*_1,*m*_1__, …, *Y*_*n*,1_ …, *Y_n,m_n__* denote the outcome vector of length *N*, where 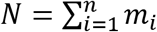 is the total number of observations. We focus on the problem of testing time-varying genetic effect between a genetic variant and the longitudinal binary trait. Let *G* denote the genotype vector of the *n* subjects at the variant to be tested, where *G_i_*, = 0, 1 or 2, depending on whether subject *i* has 0, 1 or 2 copies of minor allele at the variant.

### 2.1 GLMM with varying coefficients

We consider a generalized linear mixed model (GLMM) with varying coefficients, specified as

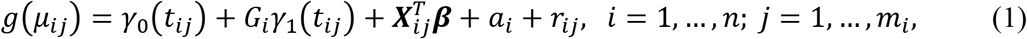

where *μ_ij_* = *E*(*Y_ij_* |*G_i_,**X**_ij_*, *α_i_*, *r_iy_*) is the mean of the response *Y_ij_* at time *t_ij_* for subject *i*, given his/her genotype, covariates, and random effects *α_i_* and *r_ij_*, *γ*_0_(*t*) and *y*_1_(*t*) are smooth nonparametric functions of time *t* representing a time-varying intercept and a time-varying genetic effect of the tested variant, ***β*** is the effects of the covariates, and *g*(*·*) is the link function. For binary traits, we use the logit link function. The correlations among repeated measurements are captured by two random effects: *a_i_* is the subject random effect and *r_ij_* is the subject-specific time-dependent random effect (Wang et al., 2017; Wu et al., 2019). We assume that *a_i_* are independent and 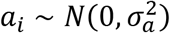. The vector of time-dependent random effects ***r**_i_* = (*r*_*i*:1_, *r_i,m_i__*) is assumed to follow a multivariate normal distribution, 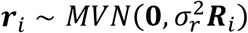, where the correlation matrix ***R**_i_* is modeled by an AR(1) structure in which *τ* is the unknown parameter. Given the random effects *a_i_* and *r_ij_*, the response *Y_ij_* are assumed to be independent. When both functions *γ*_0_(*t*) and *y*_1_(*t*) are constants, Model (1) reduces to a standard GLMM in (Wu et al., 2019).

Following (Lin and Zhang, 1999; Zhang, 2004), we estimate *γ*_0_(*t*) and *y*_1_(*t*) by maximizing the following double penalized quasi-likelihood (DPQL) function:

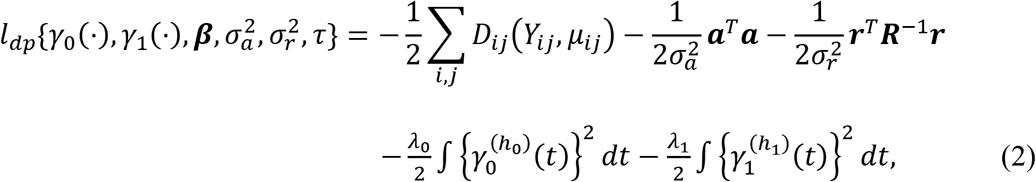

where ***a*** = (*a*_1_,…, *a_n_*)^*T*^, ***r*** = (***r***_1_…, ***r**_n_*)^*T*^ are the two vectors of random effects, ***R*** = diag{***R***_1_, _,***R**_n_*} is a block diagonal matrix, 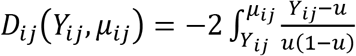 is the conditional deviance function of binary outcome *Y_ij_* given random effects *a_i_* and *r_ij_*, *λ_k_* (*k* = 0,1) are tuning parameters that control the smoothness of *γ_k_* (*t*), and *h_k_* are positive integers for the derivative order of *γ_k_* (*t*).

The maximizers for the nonparametric functions *γ_k_* (*t*) in the DPQL function of Eq. (2) are smoothing splines of order 2*h_k_* (Wahba, 1990). Let 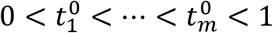 be the *m* distinct knots of *t_ij_*, the smoothing splines can be expressed as

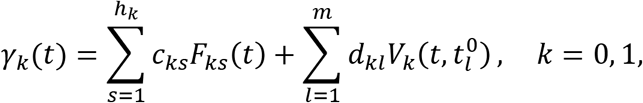

where *F_ks_* (*t*) is a polynomial of order *s* – 1 (e.g., *F_ks_*(*t*) = *t*^(*s*-1)^/(*s* – 1)!, *s* = 1 *h_k_*), and 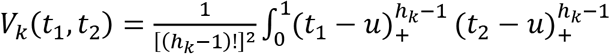 with *u*_+_ = max{*u*, 0}. We denote 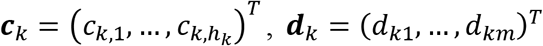 and 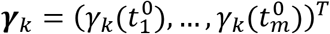 for *k* = ^0,1^.

Then ***γ**_k_* can be expressed as

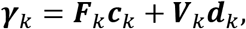

where ***F**_k_* is an *m* × *h_k_* matrix with its (*l, s*)th entry equal to 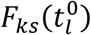, and ***V**_k_* is a positive definite matrix with the (*I, s*)th entry equal to 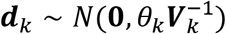. Similar to Eq. (5) in (Zhang, 2004), the DPQL function of Eq. (2) becomes

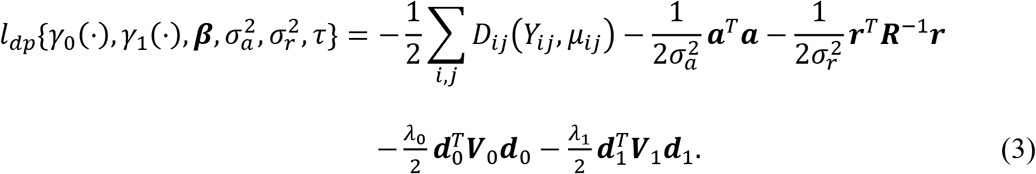

If we treat ***d**_k_* in ***γ**_k_* as random effects distributed as 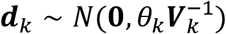 for *k* = 0,1, where 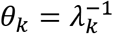, it follows that the maximizers of Eq. (3) can be obtained by fitting the GLMM representation of Model (1), expressed in a matrix form as

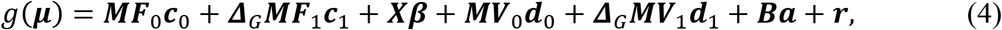

where ***M*** is an *N* × *m* incidence matrix mapping *t_ij_* to the *m* distinct knots 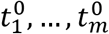, ***B*** is an *N* × *n* design matrix mapping the subject-level genotype vector ***G*** to a measurement-level genotype vector ***BG***, with its (*I, i*)th entry *B_li_* = 1 if the *l*th entry of ***Y*** is a measurement on subject *i* and 0 otherwise, and ***Δ**_G_* = diag{***BG***} = diag{*G*_1_,…, *G*_1_,…, *G_n_*,…, *G_n_*} is an *N* - dimentional diagonal matrix of the genotypes for the *n* subjects. Model (4) is a specific implementation of the GLMM with varying coefficients in (Zhang, 2004). Here, the tuning parameters *λ*_0_ and *λ*_1_ in the DPQL function of Eq. (2) are re-parameterized as *θ*_0_ and *θ*_1_, and treated as the unknown variance component parameters in Model (4).

### 2.2 Varying coefficient mixed model association test

To test time-varying genetic effect between the variant and the trait, we test *H*_0_: *y*_1_(*t*) = 0 in Model (1), which is equivalent to test *H*_0_: *c*_1_ = **0** and *θ*_1_ = 0 in Model (4). The reduced GLMM under the null hypothesis specifies that

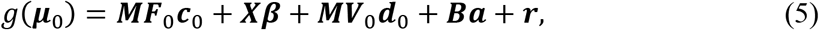

where ***μ***_0_ = *E*(***Y*** | ***F***_0_, ***X, d***_0_, ***a, r***).

If we test *H*_0_: *c*_1_ = 0 under the assumption that *θ*_1_ = 0, a score test can be constructed as

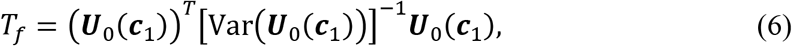

where 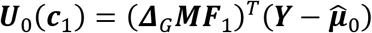 is the score function for *c*_1_ and 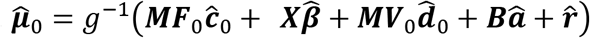 is a vector of fitted values under Model (5), which can be obtained using the penalized quasi-likelihood method (Breslow and Clayton, 1993). For binary traits, we used a bias correction procedure (Lin and Breslow, 1996; Lin and Zhang, 1999; Zhang, 2004) to produce less biased variance component estimates. Given the genotype and covariates, the variance of the score ***U***_0_(***c***_1_) under *H*_0_ is

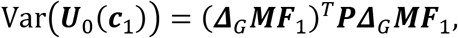

where 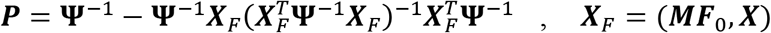 and 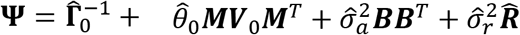. Here, 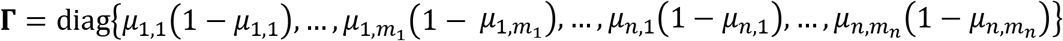 is an *N*-dimensional diagonal matrix, and 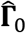 and 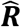 are **Γ** and ***R*** evaluated under Model (5). Under the null hypothesis, the *T_f_* test statistic has an asymptotic *χ*^2^ distribution with *h*_1_ degrees of freedom.

On the other hand, if we test *H*_0_: *θ*_1_ = 0 under the assumption that ***c***_1_ = **0**, a variance component score test can be constructed as

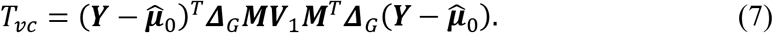

Under the null hypothesis, *T_vc_* asymptotically follows a mixture of *χ*^2^ distribution, 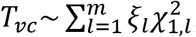, where (*ξ*_1_,…, *ξ_m_*) are the eigenvalues of the matrix 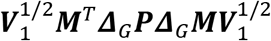 and 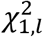 are independent 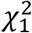 variables. The p-value of *T_vc_* can be evaluated by a moment-matching method (Liu, Tang and Zhang, 2009).

We propose a joint test for testing *H*_0_: ***c***_1_ = **0** and *θ*_1_ = 0 using a Cauchy combination test (Liu and Xie, 2020) that combines the test of fixed effect, *T_f_*, and the test of variance component, *T_vc_*, which we named as Varying-coefficient Mixed Model Association Test (VMMAT). Specifically, the VMMAT test statistic is

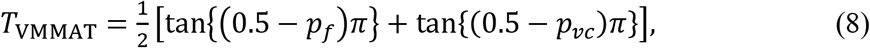

where *P_f_* and *p_vc_* are p-values of *T_f_* and *T_vc_*. Under the null hypothesis, *T*_VMMAT_ asymptotically follows a Cauchy distribution. Its p-value can be approximated by p_VMMAT_ = 0.5 — arctan (*T*_VMMAGT_)/π.

The asymptotic null distributions of *T_f_* and *T_vc_* are based on the GLMM of Eq. (4) which is an equivalent representation of the GLMM with varying coefficients using smoothing splines. Because parameters are estimated from the DPQL function of Eq. (2) with two penalty terms, the estimated variance can be larger than that from the model without penalties. Therefore, the null distributions of the score tests, *T_f_* and *T_vc_*, as well as the combined test, *T*_VMMAT_, depend on the tuning parameter values. We further assessed the null distribution of VMMAT through type I error experiments in simulation studies.

### 2.3 Retrospective varying coefficient mixed model association test

Retrospective association tests have been shown to be robust to the trait model misspecification and improve statistical power (Hayeck et al., 2015; Jiang et al., 2015; Wu et al., 2019; Wu and McPeek, 2018). In that follows, we introduce a new varying-coefficient test, RVMMAT (Retrospective Varying-coefficient Mixed Model Association Test), for testing time-varying genetic effect between the variant and the trait. RVMMAT also uses a Cauchy combination test to combine two tests: a test for *H*_0_: ***c***_1_ = 0 under the constraint *θ*_1_ = 0 and a test for *H*_0_: *θ*_1_ =0 under the constraint ***c***_1_ = 0. In contrast to the two prospective tests, *T_f_* and *T_vc_*, in VMMAT, the two tests for testing fixed effect and variance component in RVMMAT are based on a retrospective model of the genotype given the trait and covariates, such that the analysis is less dependent on the correct specification of the phenotype model. We assume that under the null hypothesis of no time-varying genetic effect between the variant and the trait, the quasi-likelihood model of the genotype *G* conditional on the phenotype *Y* and covariates *X* is

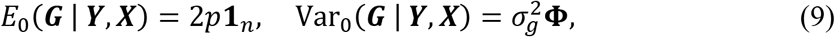

where *p* is the minor allele frequency (MAF) of the tested variant, **1**_*n*_ is a vector of length *n* with every element equals to 1, 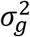 is an unknown variance parameter, and **Φ** is an *n* × *n* genetic relationship matrix (GRM) representing the overall genetic similarity between samples due to population structure, which can be estimated using genome-wide data.

When we test *H*_0_: ***c***_1_ = 0 under the assumption that *θ*_1_ = 0, the same score function ***U***_0_(***c***_1_) is considered. Because the vector of null phenotypic residuals 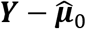, obtained by fitting Model (5), is orthogonal to the column space of ***X***_*F*_ = (***MF***_0_, ***X***), then the null mean model of ***G*** in Eq. (9) ensures that

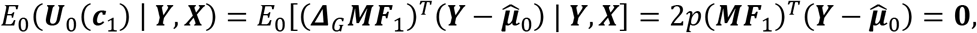

if *h*_1_ ≤ *h*_0_. In practice, we commonly use smoothing splines of the same order for *γ*_0_(*t*) and *y*_1_(*t*) in Model (1). Thus, we consider the score function ***U***_0_(***c***_1_) and construct a score test under the retrospective model of Eq. (9), given by

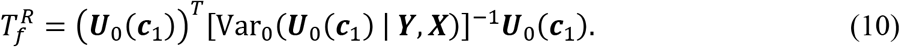

Here, the variance of ***U***_0_(***c***_1_) is evaluated by

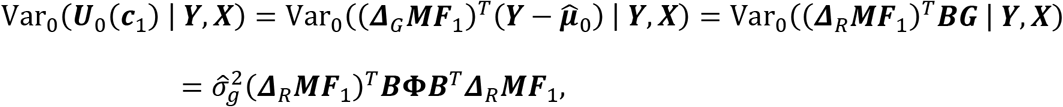

where 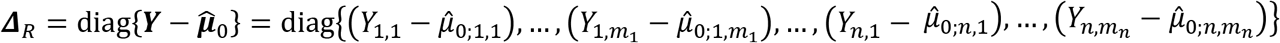 is an *N*-dimentional diagonal matrix of the phenotypic residuals. Under Hardy-Weinberg equilibrium, the variance of the genotype is estimated by 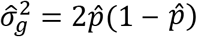, where 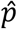 is the sample MAF of the tested variant. Under the null hypothesis, the 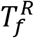 test statistic has an asymptotic *χ*^2^ distribution with *h*_1_ degrees of freedom.

If we test *H*_0_: *θ*_0_ =0 under the assumption that *c*_1_ = 0, a retrospective variance component score test under Model (9) can be constructed as

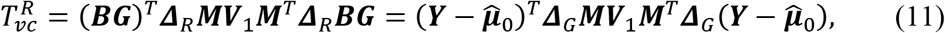

which has the same form as the prospective variance component test *T_vc_*. However, under the null hypothesis, given the trait and covariates, 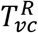 asymptotically follows a mixture of *χ*^2^ distribution, 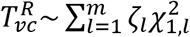, where (*ζ*_1_,…, *ζ_m_*) are the eigenvalues of the matrix 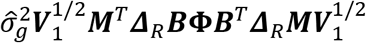.

The RVMMAT test statistic is defined by combining the two retrospective tests, 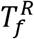 and 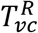, expressed as

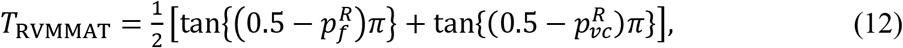

where 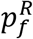 and 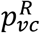 are p-values of 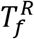 and 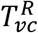. Under the null hypothesis, *T*_RVMMAT_ asymptotically follows a Cauchy distribution.

The asymptotic null distributions of 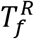 and 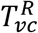 are based on the retrospective model of Eq. (9) in which genotypes are treated as random conditional on the phenotype and covariates. Therefore, the estimated variance in the trait model due to the tuning parameters in penalty terms does not impact the null distributions of the retrospective score tests, 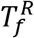 and 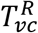, as well as the combined test, *T*_RVMMAT_. We also assessed the null distribution of RVMMAT through type I error experiments in simulation studies.

## 3 SIMULATION STUDIES

We conducted simulation studies to assess the type I error and power of VMMAT and RVMMAT, and compared them to a Gaussian copula method with weighted scores that allows for heterogenous genetic effect (Nikoloulopoulos, Joe and Chaganty, 2011) and the two association tests that assume constant genetic effect, GMMAT (Chen et al., 2016) and RGMMAT (Wu et al., 2019). In all simulations, VMMAT and RVMMAT were implemented with cubic smoothing splines. The Gaussian copula method was implemented with a binomial marginal model with the logit link function and an AR(1) structure in the Gaussian copula correlation matrix (Nikoloulopoulos et al., 2011). VMMAT and RVMMAT are designed to detect time-varying genetic effect between a genetic variant and the longitudinal binary trait. Because we test one variant at a time, these methods tend to have limited power for rare variants and are more appropriate for common variants. The performance of all methods was evaluated on common variants in simulation studies. We considered two trait models and three ascertainment schemes to evaluate the robustness of VMMAT and RVMMAT in the presence of model misspecification and ascertainment.

### 3.1 Simulation settings

To generate genotypes, we first simulated 10,000 chromosomes over a 1 Mb region using a coalescent model to mimic the recombination rates and linkage disequilibrium (LD) pattern of the European population (Schaffner et al., 2005; Shlyakhter, Sabeti and Schaffner, 2014). We then randomly selected 1,000 non-causal SNPs with MAF > 0.05. In addition, we simulated two causal SNPs that were assumed to influence the trait value with epistasis. In each simulation setting, we generated 1,000 sets of binary phenotypes at five time points for a given sample size. In the type I error experiments, for each phenotype dataset, we tested the time-varying genetic effect at the 1,000 non-causal SNPs. In total, 10^6^ test results across 1,000 phenotype datasets were used for the type I error assessment. In the power simulations, we tested time-varying genetic effect at the first of the two causal SNPs and evaluated power using 1,000 simulated phenotype datasets. In all tests considered, the genotypes at the untested SNPs were not included as covariates in the model.

We simulated binary phenotypes under two types of trait models at five time points, in which the two unlinked causal SNPs were assumed to influence phenotype through an epistatic interaction. The first type is a logistic mixed model, specified by:

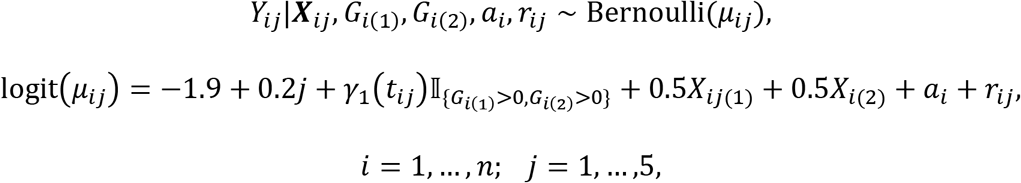

where *γ*_1_(*t_ij_*) is a function encoding the effect of the causal SNPs, *G*_*i*(1)_ and *G*_*i*(2)_ are the genotypes of subject *i* at the two causal SNPs, 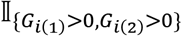 is an indicator function that takes value 1 when subject *i* has at least one copy of the minor allele at both causal SNPs, *X*_*ij*(1)_ is a continuous, time-varying covariate generated from a multivariant normal distribution with a compound symmetry correlation matrix where the correlation is 0.5, *X*_*i*(2)_ is a binary, time-invariant covariate taking values 0 or 1 with a probability of 0.5, *a_i_* and *r_ij_* are the subject-level time-independent and time-dependent random effects, respectively. We assumed 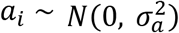 and 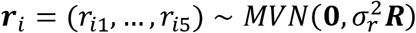, where *R* is a 5 × 5 correlation matrix specified by the AR(1) structure with a correlation coefficient *τ*. The two causal SNPs were assumed to be unlinked with MAFs 0.1 and 0.5, respectively. The variance components were set to 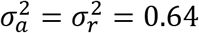 0.64 and *τ* = 0.7.

The second type of trait model is a liability threshold model in which an underlying continuous liability determines the binary outcome value based on a threshold. Specifically, the phenotype *Y_ij_* is determined by

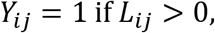

with 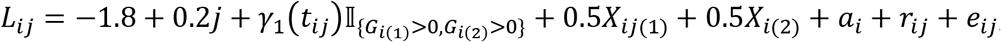, where *L_ij_* is the underlying liability for subject *i* at time *t_ij_*, and 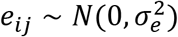 represents independent noise, with 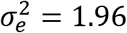. All other parameters are the same as those in the logistic mixed model.

In both trait models, we specified the intercept as a linear function of time *t_ij_* = *j*, and the genetic effect as a logistic function 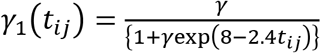 (Gong and Zou, 2012). For the type I error assessment, the effect of the causal SNPs was set to *γ* = 0.6 in *γ*_1_(*t_ij_*). For the power evaluation, we considered a range of values for *γ*, where *γ* = 0.6,0.63,0.66, and 0.69. At the given parameter values, the prevalence of the event of interest ranges from 23.68% to 40.56% over time. The proportion of the phenotypic variance explained by the two causal SNPs ranges from 0.01% to 2.99% in the logistic mixed model and from 0.01% to 1.36% in the liability threshold model.

We considered three sampling designs as in (Wu et al., 2019). In the “random” sampling, samples contain 2,000 subjects randomly selected from the population regardless of their phenotypes. In the “baseline” sampling, samples contain 1,000 case subjects and 1,000 control subjects based on their outcome value at baseline only. In the “sum” sampling, subjects were stratified into three strata based on the total count of events of the subject over time, where subjects in stratum 1 never experienced the event of interest, i.e., ∑_*j*_ *Y_ij_* = 0, subjects in stratum 2 sometimes experienced the event, i.e., 0 < ∑_*j*_*Y_ij_* < *n_i_*, and subjects in stratum 3 always experienced the event, i.e., ∑_*j*_ *Y_ij_* = *n_i_*. We oversampled subjects with response variation over the course of the study and selected 100, 1,800, and 100 subjects from the three strata (Schildcrout et al., 2018).

### 3.2 Simulation results

To assess type I error, we tested time-varying genetic effect at unlinked and unassociated SNPs. Empirical type I error was calculated as the proportion of simulations in which the p-value of the SNP is less than the nominal level *a*, for *a* = 0.01, 0.001, and 0.0001. Table 1 gives the empirical type I error rates of RVMMAT and VMMAT, based on 10^6^ replicates, under two trait models and three sampling designs. In most simulations, the type I error of RVMMAT was within the 95% confidence interval of the nominal levels. In contrast, the type I error of VMMAT in all simulation settings was much lower than the nominal level when *α* = 0.01, 0.001, and 0.0001. It is well recognized that the DPQL approach underestimates variance components when data are sparse such as binary data (Lin and Zhang, 1999; Zhang, 2004). Even with bias correction, parameters estimated from the DPQL function with penalty terms depend on the tuning parameter values. Thus, the prospective variance of the score ***U***_0_(***c***_1_) tends to be overestimated, producing a conservative test statistic. However, the retrospective variance of the score ***U***_0_(***c***_1_) does not depend on the estimation of variance components due to the tuning parameters in penalty terms so that the test statistic is less biased. These results suggest that the retrospective RVMMAT test had much better control of type I error and was robust to trait model misspecification and ascertainment, whereas the prospective VMMAT test was overly conservative.

**Table 1.** Empirical type I error of RVMMAT and VMMAT, based on 10^6^ replicates.

| Test | Nominal Level | Logistic Mixed Model |  |  | Liability Threshold Model |  |  |
| --- | --- | --- | --- | --- | --- | --- | --- |
|  |  | Random | Baseline | Sum | Random | Baseline | Sum |
| RVMMAT | 0.01 | $9.90 \times 10^{-3}$ | $1.01 \times 10^{-2}$ | $9.97 \times 10^{-3}$ | $1.02 \times 10^{-2}$ | **$9.70 \times 10^{-3}$** | $1.00 \times 10^{-2}$ |
| | 0.001 | $9.52 \times 10^{-4}$ | $9.92 \times 10^{-4}$ | **$9.17 \times 10^{-4}$** | $1.04 \times 10^{-3}$ | $1.00 \times 10^{-3}$ | $1.03 \times 10^{-3}$ |
| | 0.0001 | $9.80 \times 10^{-5}$ | $1.08 \times 10^{-4}$ | $9.40 \times 10^{-5}$ | $1.00 \times 10^{-4}$ | $1.01 \times 10^{-4}$ | $1.13 \times 10^{-4}$ |
| VMMAT | 0.01 | **$5.77 \times 10^{-3}$** | **$6.45 \times 10^{-3}$** | **$6.91 \times 10^{-3}$** | **$5.73 \times 10^{-3}$** | **$6.26 \times 10^{-3}$** | **$7.20 \times 10^{-3}$** |
|  | 0.001 | **$4.34 \times 10^{-4}$** | **$5.14 \times 10^{-4}$** | **$5.33 \times 10^{-4}$** | **$4.73 \times 10^{-4}$** | **$5.10 \times 10^{-4}$** | **$6.68 \times 10^{-4}$** |
|  | 0.0001 | **$3.70 \times 10^{-5}$** | **$4.40 \times 10^{-5}$** | **$5.70 \times 10^{-5}$** | **$2.90 \times 10^{-5}$** | **$5.00 \times 10^{-5}$** | **$4.60 \times 10^{-5}$** |

Rates outside of the 95% confidence interval are in bold.

To compare power, we considered four parameter values for *γ* to determine time-varying genetic effect at the two causal SNPs and tested between the trait and the first causal SNP.

Empirical power was calculated at the significance level 10^−3^, based on 1,000 replicates. Figure 1 demonstrates the power results of the five methods, RVMMAT, VMMAT, Copula, RGMMAT and GMMAT, under two trait models and three sampling designs. In all simulation settings, the two varying-coefficient tests consistently had higher power than the association tests assuming constant gene effect. The Gaussian copula method with heterogenous genetic effect had lower power than RVMMAT and VMMAT, while performed better than RGMMAT and GMMAT. Moreover, within the same type of tests, the retrospective test was more powerful than the prospective test. Both RVMMAT and VMMAT had similar power across the three sampling designs. In contrast, Copula, RGMMAT and GMMAT had lower power under the sum sampling in both trait models. The power gain of the varying-coefficient tests was more prominent over the association tests assuming constant genetic effect in the presence of ascertainment. These results suggest that RVMMAT was the most powerful test and outperformed the association tests assuming constant genetic effect.

**Figure 1.**
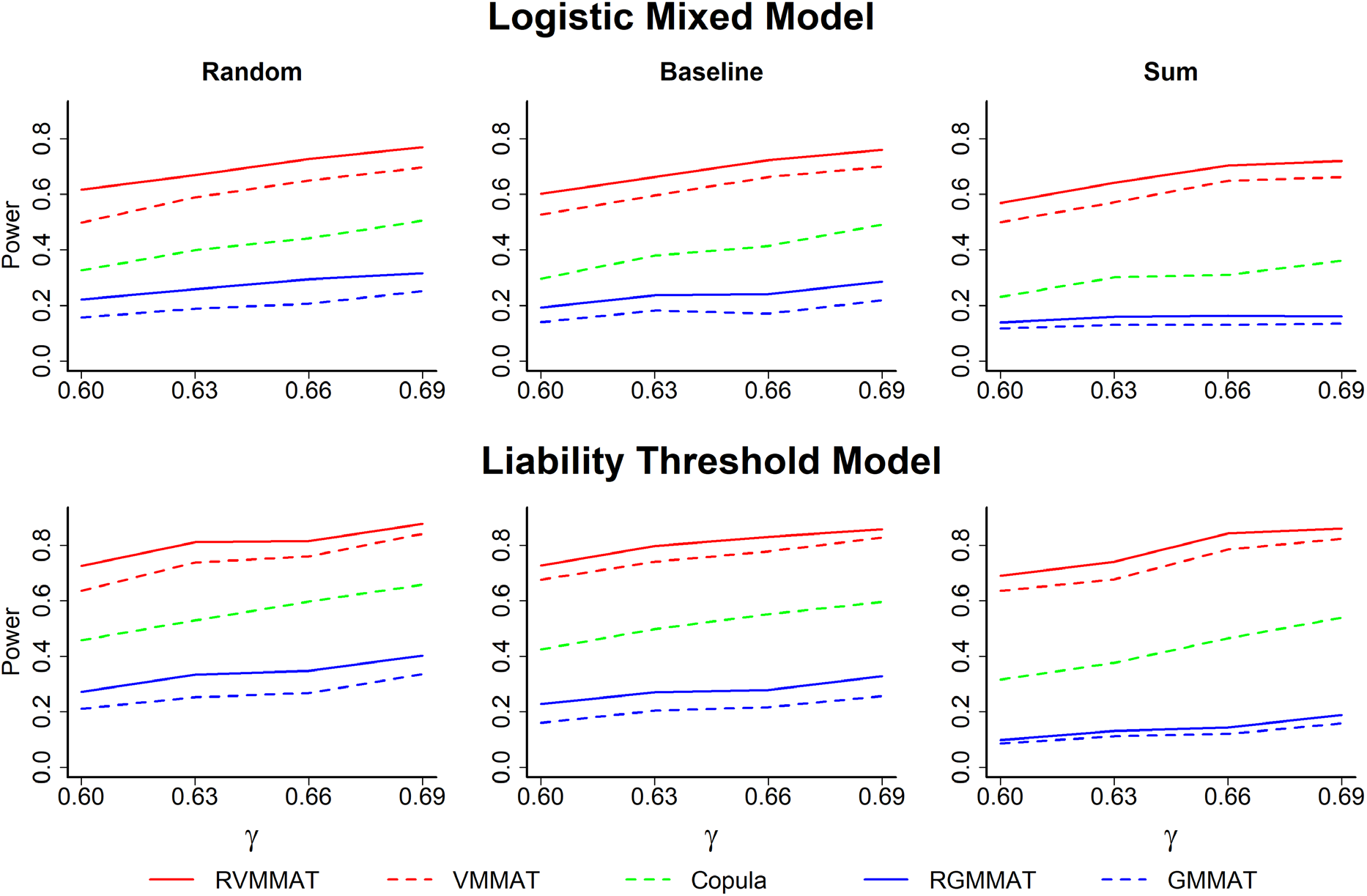
Empirical power of RVMMAT, VMMAT, Copula, RGMMAT and GMMAT. Power is based on 1,000 replicates at five time points with *α* = 10^−3^. In the upper panel, trait is simulated under the logistic mixed model; in the lower panel, trait is simulated under the liability threshold model. Power results are demonstrated in samples of 2,000 individuals according to three ascertainment schemes: random, baseline, and sum.

## 4 APPLICATION TO MESA DATA

We applied our proposed methods to a genome-wide association analysis of hypertension in MESA (Bild et al., 2002). MESA is a large longitudinal study of subclinical cardiovascular disease (CVD) whose primary objective is to understand the pathogenesis of atherosclerosis and other CVD. We analyzed longitudinal hypertension assessed at five time points on 6,429 participants. Among them, 39.3% are white, 26.1% are African American, 22.5% are Hispanic, and 12.1% are Asian. The proportion of case subjects at each time point ranges from 44.6% (*n* = 2,864) to 59.5% (*n* = 2,608), and the missing rate at each time point ranges from 0 to 31.6%.

Samples were genotyped using the Affymetrix Human SNP Array 6.0. After data cleaning, there were 6,428 subjects available for genotype imputation. We applied IMPUTE2 (Howie, Donnelly and Marchini, 2009) for imputation, using the 1000 Genomes Phase 3 data as a reference panel. Subjects who did not meet either of the following criteria were excluded: (1) proportion of successfully imputed SNPs > 95% and (2) empirical inbreeding coefficient < 0.05. Based on the above criteria, 6,424 subjects were retained in the downstream analysis, with 3,057 males and 3,367 females, of whom 2,527 are white, 1,673 are African American, 1,449 are Hispanic, and 775 are Asian. There are 2,227 subjects who had no hypertension during the study period, 1,807 subjects who were sometimes hypertensive, i.e., exhibited response variation, and 2,390 subjects who were always hypertensive over the course of the study. We then tested Hardy-Weinberg equilibrium at each SNP within each population. SNPs met all of the following quality-control conditions were included in the analysis: (1) call rate > 95%, (2) Hardy-Weinberg *χ*^2^ statistic p-value > 10^−6^, and (3) MAF > 1%. Taken together, a final set of 6,155,404 SNPs were examined in the downstream analysis.

### 4.1 Analysis of time-varying genetic effect

We performed genome-wide tests of time-varying genetic effect on hypertension using RVMMAT and VMMAT with cubic smoothing splines in the MESA sample. Age at baseline, sex, and the top ten principal components (PCs) were included as time-invariant covariates in the analysis. The top ten PCs were calculated using the LD pruned SNPs with MAF > 0.05 to control for population structure. Since hypertension was assessed in year 2000, 2002, 2004, 2005 and 2010, we coded time at the five time points as 0, 0.2, 0.4, 0.5 and 1, respectively. We also applied the Gaussian copula method with heterogenous genetic effect, adjusting for the same covariates. To compare the performance of the varying-coefficient tests with the association tests assuming constant genetic effect, we applied RGMMAT and GMMAT to the analysis of hypertension, adjusting for age at baseline, sex, time, and the top ten PCs.

The two retrospective tests, RVMMAT and RGMMAT, showed no evidence of inflation in the quantile-quantile (Q-Q) plot. The genomic control inflation factors were 0.905 and 0.976, respectively. The prospective VMMAT test was overly conservative, with a genomic control factor of 0.774, consistent with the observed deflation in the type I error simulations. The genomic control inflation factor was 0.838 for GMMAT.

None of the SNPs reached genome-wide significance at the p-value threshold of 5 × 10^−8^ that is widely used in GWAS. Table 2 reports the top SNPs for which at least one of the tests gives a p-value < 5 × 10^−7^. The smallest p-values of these eight SNPs were mostly generated by RVMMAT, except at the last two SNPs. VMMAT generated much larger p-values than RVMMAT due to its conservativeness, while RGMMAT and GMMAT had comparable results. The Gaussian copula method produced p-values comparable to VMMAT, except at the last two SNPs. A cluster of six SNPs in LD (*r*^2^ > 0.97), rs145659245, rs58265184, rs57719815, rs60197637, rs61327798, and rs142890225, located at 4p15, showed time-varying genetic effect on hypertension by RVMMAT (p-value = 6.78 × 10^−8^ — 2.85 × 10^−7^). Figure 2A demonstrated the estimated genetic effect over time at these SNPs where the estimated effect at each time point was obtained by using the observed trait values at that time point only. A strait line was used to connect the estimated values at two adjacent time points. We observed an increasing and then decreasing trend in genetic effects on hypertension across the five time points. However, RGMMAT and GMMAT lost power and generated large p-values by assuming constant genetic effect. These SNPs are in an intron of the gene *PROM1*, encoding a pentaspan transmembrane glycoprotein, which was reported to be associated with pulse pressure (Evangelou et al., 2018). The smallest p-value for rs374012917, located on chromosome 17, was generated by the Gaussian copula method (p-value = 1.02 × 10^−7^). As the estimated genetic effects at the five time points were relatively stable for this SNP (Figure 2B), RGMMAT and GMMAT generated slightly larger p-values. There was also evidence of association between hypertension and rs72930733 (p-value = 2.55 × 10^−7^). Although RVMMAT did not give the smallest p-value for this SNP, its p-value was slightly larger than that of RGMMAT, mostly due to the increasing trend in genetic effect (Figure 2B). This SNP is in an intron of the gene *WDR7*, located at 18q21. Two hypertension GWAS identified an association between *WDR7* and systolic blood pressure (Evangelou et al., 2018; Kichaev et al., 2019).

**Figure 2.**
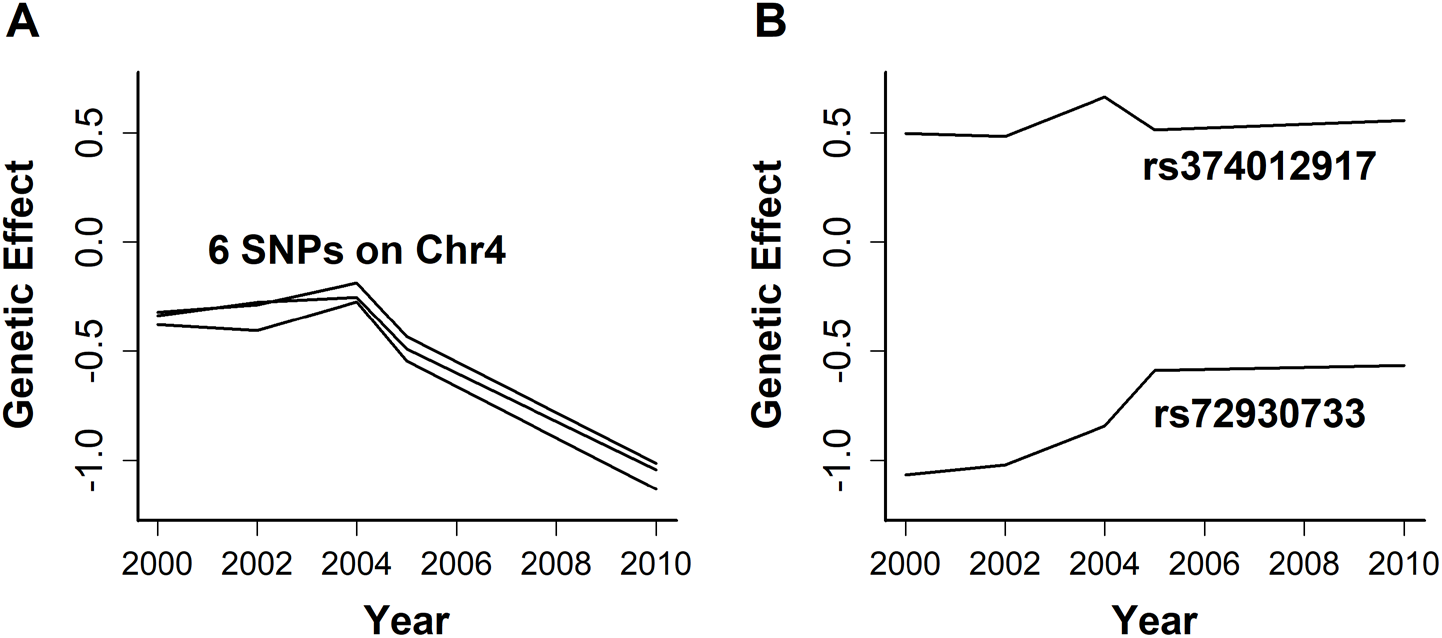
Estimated genetic effect of the top 8 SNPs on hypertension at each of the five time points. (A) six SNPs on chromosome 4; (B) two SNPs on chromosomes 17 and 18, respectively.

**Table 2.** SNPs with p-value < 5×10 ^7^ in at least one of the tests in the MESA data.

| Chr | Gene Region | SNP | Position | MAF | RVMMAT | VMMAT | Copula | RGMMAT | GMMAT |
| --- | --- | --- | --- | --- | --- | --- | --- | --- | --- |
| 4 | *PROM1* | rs145659245 | 16,060,553 | 0.013 | **$6.78 \times 10^{-8}$** | $4.19 \times 10^{-6}$ | $9.77 \times 10^{-6}$ | $7.67 \times 10^{-4}$ | $1.78 \times 10^{-3}$ |
| | | rs58265184 | 16,061,151 | 0.014 | **$2.18 \times 10^{-7}$** | $7.70 \times 10^{-5}$ | $2.79 \times 10^{-5}$ | $2.51 \times 10^{-3}$ | $4.85 \times 10^{-3}$ |
| | | rs57719815 | 16,063,652 | 0.013 | **$2.85 \times 10^{-7}$** | $8.04 \times 10^{-5}$ | $3.09 \times 10^{-5}$ | $1.65 \times 10^{-3}$ | $3.27 \times 10^{-3}$ |
| | | rs60197637 | 16,063,659 | 0.013 | **$2.85 \times 10^{-7}$** | $8.04 \times 10^{-5}$ | $3.09 \times 10^{-5}$ | $1.65 \times 10^{-3}$ | $3.27 \times 10^{-3}$ |
| | | rs61327798 | 16,063,661 | 0.013 | **$2.85 \times 10^{-7}$** | $8.04 \times 10^{-5}$ | $3.09 \times 10^{-5}$ | $1.65 \times 10^{-3}$ | $3.27 \times 10^{-3}$ |
| | | rs142890225 | 16,065,544 | 0.013 | **$2.85 \times 10^{-7}$** | $8.04 \times 10^{-5}$ | $3.10 \times 10^{-5}$ | $1.65 \times 10^{-3}$ | $3.27 \times 10^{-3}$ |
| 17 | *LRRC37B* | rs374012917 | 30,403,054 | 0.038 | $2.21 \times 10^{-5}$ | $1.95 \times 10^{-4}$ | **$1.02 \times 10^{-7}$** | $2.31 \times 10^{-6}$ | $4.95 \times 10^{-6}$ |
| 18 | *WDR7* | rs72930733 | 54,641,870 | 0.011 | $1.80 \times 10^{-6}$ | $7.06 \times 10^{-5}$ | $7.72 \times 10^{-6}$ | **$2.55 \times 10^{-7}$** | $4.63 \times 10^{-6}$ |

The smallest p-values among the five tests at the given SNPs are in bold.

We further assessed the model fitting of cubic smoothing splines on the top SNPs in Table 2 using deviance and goodness-of-fit p-value. All p-values were large, suggesting that there was no evidence of lack of fit (Table 3). We also checked the deviance residuals of the cubic smoothing splines model applied to the top SNPs. The deviance residuals range from −2.46 to 2.19, suggesting that cubic smoothing splines fit the data adequately.

**Table 3.** Assessment of model fitting with cubic smoothing splines at the top SNPs in the MESA data.

| Chr | Gene Region | SNP | Position | MAF | Deviance | Goodness-of-fit<br>P-value |
| --- | --- | --- | --- | --- | --- | --- |
| 4 | *PROM1* | rs145659245 | 16,060,553 | 0.013 | 10467.93 | 1 |
|  |  | rs58265184 | 16,061,151 | 0.014 | 10468.13 | 1 |
|  |  | rs57719815 | 16,063,652 | 0.013 | 10469.79 | 1 |
|  |  | rs60197637 | 16,063,659 | 0.013 | 10469.79 | 1 |
|  |  | rs61327798 | 16,063,661 | 0.013 | 10469.79 | 1 |
|  |  | rs142890225 | 16,065,544 | 0.013 | 10469.79 | 1 |
| 17 | *LRRC37B* | rs374012917 | 30,403,054 | 0.038 | 10495.83 | 1 |
| 18 | *WDR7* | rs72930733 | 54,641,870 | 0.011 | 10493.05 | 1 |

### 4.2 Pathway analysis

We then performed functional pathway analysis using the MetaCore^TM^ software to identify enriched pathways related to hypertension. The top SNPs for which at least one of the tests had a p-value < 2 × 10^−4^ were included in the analysis. Fisher’s exact test was used to determine whether the SNP list was enriched for a functional pathway. At the false discovery rate (FDR) < 0.05, we identified two significant pathways that were associated with G-protein signaling and DNA damage. The first one is the G-protein signaling pathway related to Rac1 activation (p-value = 6.12 × 10^−5^, FDR = 1.65 × 10^−2^). Rac1 participates in the control of blood pressure through multiple mechanisms in the arterial wall and the central nervous system (Loirand and Pacaud, 2010). Importantly, a role for Rac1 in atherosclerosis and cardiac hypertrophy has been established in response to the administration of statins in clinical trials (Maack et al., 2003). Animal studies indicated that Rac1 is essential for endothelium-dependent vasomotor response, the redox state of blood vessels and homeostasis of blood pressure (Moustafa-Bayoumi et al., 2003; Satoh et al., 2006; Sawada et al., 2008). The second pathway is the DNA damage pathway related to the ataxia-telangiectasia mutated (ATM) kinase activation (p-value = 4.98 × 10^−4^, FDR = 4.48 × 10^−2^). Emerging evidence has demonstrated that accumulated DNA damage and subsequent repair pathways play a crucial role in the initiation and progression of cardiovascular disorders, such as atherosclerosis and maladaptive cardiac hypertrophy (Shah et al., 2018; Shah and Mahmoudi, 2015; Uryga, Gray and Bennett, 2016; Wu et al., 2022). ATM-mediated phosphorylation plays cardinal roles in response to genomic stress to preserve cellular homeostasis. DNA double-strand breaks trigger ATM activation which mediates DNA damage response and regulate cardiac remodeling, inflammation, and systolic function, eventually promoting heart failure development (Shiloh and Ziv, 2013; Uziel et al., 2003).

## 5 DISCUSSION

In genome-wide association analysis of longitudinal traits, modeling time-varying genetic effect can increase power for the detection of genes underlying the development and progression of complex diseases. In this study, we developed RVMMAT, a GLMM-based, retrospective varying-coefficient association testing method for longitudinal binary traits. RVMMAT extends the existing association methods assuming constant effect over time to testing of time-varying effect on binary traits. RVMMAT is constructed based on the trait model allowing for time-varying genetic effect. The variance of the test statistics is assessed retrospectively by considering the conditional distribution of the genotype at the variant of interest, given phenotype and covariate information, under the null hypothesis of no association. RVMMAT has the following features: (1) it is computationally feasible for genetic studies with millions of variants, (2) it has well-controlled type I error in the presence of ascertainment and trait model misspecification, and (3) it can easily be fitted as a GLMM model using popular software such as R and SAS. We also propose VMMAT, a prospective varying-coefficient association test, for performance comparison.

Our simulation results demonstrated that RVMMAT maintained correct type I error under different trait models and ascertainment schemes, whereas VMMAT was overly conservative due to the biased estimation of variance in the penalized trait model. We further demonstrated that the retrospective RVMMAT test achieved the highest power among the five tests under all the trait models and ascertainment schemes considered in the simulations. Application of RVMMAT to the MESA longitudinal hypertension data identified three novel genes that were associated with hypertension. Among them, two genes are known to be associated with systolic blood pressure and pulse pressure. Moreover, we identified two significant pathways associated with longitudinal hypertension: the G-protein signaling pathway related to Rac1 activation, and the DNA damage pathway related to ATM activation. Given the established role for Rac1 and ATM in atherosclerosis and cardiac hypertrophy, our findings suggest that RVMMAT can provide enhanced statistical power in detecting biologically relevant genetic loci that are associated with trait dynamics. A better understanding of temporal variation of trait values and time-varying genetic contribution may shed light on the genetic mechanisms influencing the temporal trend of diseases and complex traits.

The RVMMAT and VMMAT methods are designed for single-variant association analysis of longitudinal binary traits. However, single-variant association tests suffer from restricted power to detect association for rare variants in whole-genome sequencing studies. As many variants influence complex traits collectively, assessing joint effects from multiple variants by aggregating weak signals at the gene or pathway level holds great promise for the identification of novel genes underlying disease risks. To extend RVMMAT to rare variant analysis with longitudinal binary data, we could design a linear statistic or a quadratic statistic that combines the test allowing for time-varying genetic effect at each variant in a gene region. Such statistics are likely to better calibrate the fluctuation of genetic contributions to the trait values over time.

Additionally, our current model can easily be extended to analyze nominal, ordinal and count data.

## ACKNOWLEDGMENTS

The authors would like to thank the reviewers, Associate Editor and Editor for their feedback that substantially improved the quality of this paper.

## SUPPLEMENTARY MATERIAL

R package implementing RVMMAT and VMMAT and simulation code can be found at https://github.com/ZWang-Lab/RVMMAT.

## REFERENCES

Bild, D. E., Bluemke, D. A., Burke, G. L., et al. (2002). Multi-ethnic study of atherosclerosis: objectives and design. American journal of epidemiology 156, 871–881.

Breslow, N. E., and Clayton, D. G. (1993). Approximate Inference in Generalized Linear Mixed Models. Journal of the American Statistical Association 88, 9–25.

Bryois, J., Buil, A., Ferreira, P. G., et al. (2017). Time-dependent genetic effects on gene expression implicate aging processes. Genome research 27, 545–552.

Chen, H., Wang, C., Matthew, et al. (2016). Control for Population Structure and Relatedness for Binary Traits in Genetic Association Studies via Logistic Mixed Models. The American Journal of Human Genetics 98, 653–666.

Chiang, C.-T., Rice, J. A., and Wu, C. O. (2001). Smoothing spline estimation for varying coefficient models with repeatedly measured dependent variables. Journal of the American Statistical Association 96, 605–619.

Chu, W., Li, R., Liu, J., and Reimherr, M. (2020). Feature selection for generalized varying coefficient mixed-effect models with application to obesity GWAS. The Annals of Applied Statistics 14, 276–298.

Chu, W., Li, R., and Reimherr, M. (2016). Feature screening for time-varying coefficient models with ultrahigh dimensional longitudinal data. The Annals of Applied Statistics 10, 596.

Das, K., Li, J., Wang, Z, et al. (2011). A dynamic model for genome-wide association studies. Human genetics 129, 629–639.

Eubank, R., Huang, C., Maldonado, Y. M., Wang, N., Wang, S., and Buchanan, R. (2004). Smoothing spline estimation in varying-coefficient models. Journal of the Royal Statistical Society: Series B (Statistical Methodology) 66, 653–667.

Evangelou, E., Warren, H. R., Mosen-Ansorena, D., et al. (2018). Genetic analysis of over 1 million people identifies 535 new loci associated with blood pressure traits. Nature Genetics 50, 1412–1425.

Fan, J., Ma, Y., and Dai, W. (2014). Nonparametric independence screening in sparse ultra-high-dimensional varying coefficient models. Journal of the American Statistical Association 109, 1270–1284.

Fan, J., and Zhang, W. (1999). Statistical estimation in varying coefficient models. The annals of Statistics 27, 1491–1518.

Fan, J., and Zhang, W. (2008). Statistical methods with varying coefficient models. Statistics and its Interface 1, 179.

Furlotte, N. A., Eskin, E., and Eyheramendy, S. (2012). Genome-wide association mapping with longitudinal data. Genetic epidemiology 36, 463–471.

Gong, Y., and Zou, F. (2012). Varying coefficient models for mapping quantitative trait loci using recombinant inbred intercrosses. Genetics 190, 475–486.

Hastie, T., and Tibshirani, R. (1993). Varying - coefficient models. Journal of the Royal Statistical Society: Series B (Statistical Methodology) 55, 757–779.

Hayeck, T. J., Zaitlen, N. A., Loh, P.-R., et al. (2015). Mixed model with correction for casecontrol ascertainment increases association power. The American Journal of Human Genetics 96, 720–730.

Hoover, D. R., Rice, J. A., Wu, C. O., and Yang, L.-P. (1998). Nonparametric smoothing estimates of time-varying coefficient models with longitudinal data. Biometrika 85, 809–822.

Howie, B. N., Donnelly, P., and Marchini, J. (2009). A flexible and accurate genotype imputation method for the next generation of genome-wide association studies. PLoS genetics 5, e1000529.

Huang, J. Z., and Shen, H. (2004). Functional coefficient regression models for non-linear time series: a polynomial spline approach. Scandinavian journal of statistics 31, 515–534.

Huang, J. Z., Wu, C. O., and Zhou, L. (2002). Varying-coefficient models and basis function approximations for the analysis of repeated measurements. Biometrika 89, 111–128.

Huang, J. Z., Wu, C. O., and Zhou, L. (2004). Polynomial spline estimation and inference for varying coefficient models with longitudinal data. Statistica Sinica, 763–788.

Jiang, D., Mbatchou, J., and McPeek, M. S. (2015). Retrospective association analysis of binary traits: overcoming some limitations of the additive polygenic model. Human heredity 80, 187–195.

Joe, H. (2014). Dependence modeling with copulas: CRC press.

Kauermann, G., and Tutz, G. (1999). On model diagnostics using varying coefficient models. Biometrika 86, 119–128.

Kichaev, G., Bhatia, G., Loh, P. R., et al. (2019). Leveraging Polygenic Functional Enrichment to Improve GWAS Power. The American Journal of Human Genetics 104, 65–75.

Kürüm, E., Hughes, J., Li, R., and Shiffman, S. (2018). Time-varying copula models for longitudinal data. Statistics and its Interface 11, 203.

Kürüm, E., Li, R., Shiffman, S., and Yao, W. (2016). Time-varying coefficient models for joint modeling binary and continuous outcomes in longitudinal data. Statistica Sinica 26, 979.

Li, J., Wang, Z., Li, R., and Wu, R. (2015). Bayesian group Lasso for nonparametric varying-coefficient models with application to functional genome-wide association studies. The Annals of Applied Statistics 9, 640–664.

Liang, K.-Y., and Zeger, S. L. (1986). Longitudinal data analysis using generalized linear models. Biometrika 73, 13–22.

Lin, X., and Breslow, N. E. (1996). Bias correction in generalized linear mixed models with multiple components of dispersion. Journal of the American Statistical Association 91, 1007–1016.

Lin, X., and Carroll, R. J. (2000). Nonparametric function estimation for clustered data when the predictor is measured without/with error. Journal of the American Statistical Association 95, 520–534.

Lin, X., and Zhang, D. (1999). Inference in generalized additive mixed modelsby using smoothing splines. Journal of the Royal Statistical Society: Series B (Statistical Methodology) 61, 381–400.

Liu, H., Tang, Y., and Zhang, H. H. (2009). A new chi-square approximation to the distribution of non-negative definite quadratic forms in non-central normal variables. Computational Statistics & Data Analysis 53, 853–856.

Liu, J., Li, R., and Wu, R. (2014). Feature selection for varying coefficient models with ultrahigh-dimensional covariates. Journal of the American Statistical Association 109, 266–274.

Liu, Y., and Xie, J. (2020). Cauchy combination test: a powerful test with analytic p-value calculation under arbitrary dependency structures. Journal of the American Statistical Association 115, 393–402.

Loirand, G., and Pacaud, P. (2010). The role of Rho protein signaling in hypertension. Nature Reviews Cardiology 7, 637–647.

Londono, D., Chen, K.-m., Musolf, A., et al. (2013). A novel method for analyzing genetic association with longitudinal phenotypes. Statistical applications in genetics and molecular biology 12, 241–261.

Lu, Y., and Zhang, R. (2009). Smoothing spline estimation of generalised varying-coefficient mixed model. Journal of Nonparametric Statistics 21, 815–825.

Maack, C., Kartes, T., Kilter, H., et al. (2003). Oxygen free radical release in human failing myocardium is associated with increased activity of rac1-GTPase and represents a target for statin treatment. Circulation 108, 1567–1574.

Marchetti-Bowick, M., Yin, J., Howrylak, J. A., and Xing, E. P. (2016). A time-varying group sparse additive model for genome-wide association studies of dynamic complex traits. Bioinformatics 32, 2903–2910.

Meirelles, O. D., Ding, J., Tanaka, T., et al. (2013). SHAVE: shrinkage estimator measured for multiple visits increases power in GWAS of quantitative traits. European Journal of Human Genetics 21, 673–679.

Moustafa-Bayoumi, M., Wisel, S., Goldschmidt-Clermont, P. J., and Hassanain, H. H. (2003). Hypertension caused by transgenic overexpression of Rac1. Medicine & Science in Sports & Exercise 35, S186.

Nikoloulopoulos, A. K., Joe, H., and Chaganty, N. R. (2011). Weighted scores method for regression models with dependent data. Biostatistics 12, 653–665.

Ning, C., Kang, H., Zhou, L., et al. (2017). Performance Gains in Genome-Wide Association Studies for Longitudinal Traits via Modeling Time-varied effects. Scientific Reports 7.

Satoh, M., Ogita, H., Takeshita, K., Mukai, Y., Kwiatkowski, D. J., and Liao, J. K. (2006). Requirement of Rac1 in the development of cardiac hypertrophy. Proceedings of the National Academy of Sciences 103, 7432–7437.

Sawada, N., Salomone, S., Kim, H.-H., Kwiatkowski, D. J., and Liao, J. K. (2008). Regulation of endothelial nitric oxide synthase and postnatal angiogenesis by Rac1. Circulation research 103, 360–368.

Schaffner, S. F., Foo, C., Gabriel, S., Reich, D., Daly, M. J., and Altshuler, D. (2005). Calibrating a coalescent simulation of human genome sequence variation. Genome research 15, 1576–1583.

Schildcrout, J. S., Schisterman, E. F., Mercaldo, N. D., Rathouz, P. J., and Heagerty, P. J. (2018). Extending the Case-Control Design to Longitudinal Data. Epidemiology 29, 67–75.

Shah, A., Gray, K., Figg, N., Finigan, A., Starks, L., and Bennett, M. (2018). Defective base excision repair of oxidative DNA damage in vascular smooth muscle cells promotes atherosclerosis. Circulation 138, 1446–1462.

Shah, N. R., and Mahmoudi, M. (2015). The role of DNA damage and repair in atherosclerosis: A review. Journal of molecular and cellular cardiology 86, 147–157.

Shiloh, Y., and Ziv, Y. (2013). The ATM protein kinase: regulating the cellular response to genotoxic stress, and more. Nature reviews Molecular cell biology 14, 197–210.

Shlyakhter, I., Sabeti, P. C., and Schaffner, S. F. (2014). Cosi2: an efficient simulator of exact and approximate coalescent with selection. Bioinformatics 30, 3427–3429.

Sikorska, K., Rivadeneira, F., Groenen, P. J., et al. (2013). Fast linear mixed model computations for genome - wide association studies with longitudinal data. Statistics in medicine 32, 165–180.

Sitlani, C. M., Rice, K. M., Lumley, T., et al. (2015). Generalized estimating equations for genome-wide association studies using longitudinal phenotype data. Statistics in medicine 34, 118–130.

Uryga, A., Gray, K., and Bennett, M. (2016). DNA damage and repair in vascular disease. Annual review of physiology 78, 45–66.

Uziel, T., Lerenthal, Y., Moyal, L., Andegeko, Y., Mittelman, L., and Shiloh, Y. (2003). Requirement of the MRN complex for ATM activation by DNA damage. The EMBO journal 22, 5612–5621.

Wahba, G. (1990). Spline models for observational data: Society for industrial and applied mathematics.

Wang, L., Li, H., and Huang, J. Z. (2008). Variable selection in nonparametric varying-coefficient models for analysis of repeated measurements. Journal of the American Statistical Association 103, 1556–1569.

Wang, Z., Xu, K., Zhang, X., Wu, X., and Wang, Z. (2017). Longitudinal SNP-set association analysis of quantitative phenotypes. Genetic epidemiology 41, 81–93.

Wu, C. O., Chiang, C.-T., and Hoover, D. R. (1998). Asymptotic confidence regions for kernel smoothing of a varying-coefficient model with longitudinal data. Journal of the American Statistical Association 93, 1388–1402.

Wu, L., Sowers, J. R., Zhang, Y., and Ren, J. (2022). Targeting DNA damage response in cardiovascular diseases: from pathophysiology to therapeutic implications. Cardiovascular research.

Wu, W., Wang, Z., Xu, K., et al. (2019). Retrospective Association Analysis of Longitudinal Binary Traits Identifies Important Loci and Pathways in Cocaine Use. Genetics 213, 1225-1236.

Wu, X., and McPeek, M. S. (2018). L-gator: genetic association testing for a longitudinally measured quantitative trait in samples with related individuals. The American Journal of Human Genetics 102, 574–591.

Xia, X., Yang, H., and Li, J. (2016). Feature screening for generalized varying coefficient models with application to dichotomous responses. Computational Statistics & Data Analysis 102, 85–97.

Zhang, D. (2004). Generalized Linear Mixed Models with Varying Coefficients for Longitudinal Data. Biometrics 60, 8–15.

